# Internal in-frame translation generates Cas11b, which is important for effective interference in an archaeal CRISPR-Cas system

**DOI:** 10.1101/2024.10.02.616218

**Authors:** A.-L. Sailer, J. Brendel, A. Chernev, S. König, T. Bischler, T. Gräfenhan, H. Urlaub, U. Gophna, A. Marchfelder

## Abstract

CRISPR-Cas is a sophisticated defence system used by bacteria and archaea to fend off invaders. CRISPR-Cas systems vary in their Cas protein composition and have therefore been divided into different classes and types. Type I systems of bacteria have been shown to contain the small Cas11 protein as part of the interference complex. Here we show for the first time that an archaeal CRISPR-Cas type I system also contains a Cas11 protein. In addition, we show for the first time an internal in-frame translation of an archaeal protein. The Cas11b protein from the *Haloferax volcanii* type I-B system is encoded in the *cas8b* gene. Translation initiation at an internal methionine of the *cas8b* open reading frame results in synthesis of Cas11b. Cas11b is required for an effective interference reaction and without Cas11b fewer Cascade complexes form. Comparison of transcriptomes from wild type and a Cas11b less strain show that the depletion of Cas11b results in differential regulation of many genes. Taken together Cas11b is important for the defence reaction of the type I-B CRISPR-Cas system and seems to play an additional cellular role.

## Introduction

The CRISPR-Cas immune system is one of several systems evolved by bacteria and archaea to defend themselves against invaders (Hille *et al*, 2018; Koonin *et al*, 2017; Makarova *et al*, 2020). In contrast to previous defence systems the CRISPR-Cas system is special because it adapts to new viruses and makes the cell immune to recurring attacks. Many different versions of the CRISPR-Cas systems have been found and described, however only a few of them have been characterised in detail (Makarova *et al*., 2020). All CRISPR-Cas systems have in common that they employ Cas proteins and short RNAs (known as crRNAs) for the defence reaction. They all use three stages to fend off the invader: adaptation, expression and interference. The adaptation step makes the CRISPR-Cas system more sophisticated than other defence systems since it renders the cell immune against new mobile genetic elements. Upon detection of the invader its DNA is degraded and a piece of it is integrated into the CRISPR locus. The integration of the invader DNA into the genome makes the defence hereditary. In the expression step the CRISPR RNA containing the information from all previous invaders, is expressed and processed into the invader specific crRNAs, each being able to detect an invader during a recurring attack. In the last step, the interference step, the invader DNA is recognised by its specific crRNA and immediately degraded. This last step is very efficient ensuring cell survival.

One major difference between the many CRISPR-Cas systems found is the nature of the effector that detects and degrades the invader. In class 2 systems the effector is a single protein like Cas9 while class 1 systems use a multiprotein complex as effector (Makarova *et al*., 2020). There are three different class 1 types: type I, III and IV with type I systems occuring most frequently (Makarova *et al*., 2020). The major types are further subdivided into subtypes based on the nature of their Cas proteins. For type I these are subtypes I-A to I-G (Makarova *et al*., 2020), with I-B being the most abundant subtype. The effector complex of the type I systems is termed Cascade (<u>C</u>RISPR-<u>as</u>sociated <u>c</u>omplex for <u>a</u>ntiviral <u>de</u>fence).

While CRISPR-Cas systems have been found in archaea and bacteria, they are more prevalent in archaea with about 80% having this defence system, while only approximately 40% of bacteria encode a CRISPR-Cas system (Makarova *et al*., 2020). The archaeon *Haloferax volcanii* has a single CRISPR-Cas system of type I-B. The *Haloferax* system has been studied in detail and an efficient CRISPRi tool has been developed on its basis (Brendel *et al*., 2014; Fischer *et al*, 2012; Gophna *et al*, 2017; Maier *et al*, 2017; Maier *et al*, 2013; Maier *et al*, 2019; Maier *et al*, 2015; Miezner *et al*, 2023; Stachler & Marchfelder, 2016). Its Cascade complex consists of Cas5 and Cas7 as well as a loosely associated Cas8b. Cas6b was copurified with the complex, but is not required for the interference rendering it a non-integral part of the complex (Fig. 1B.) (Brendel *et al*., 2014; Maier *et al*., 2015). Cascade complexes from type I-A and type I-E systems have been shown to contain an additional small subunit, termed Cas11 or small subunit, which is encoded by a separate gene (Fig. 1A.) (Makarova *et al*., 2020). Cas11 proteins from the different subtypes do not show significant sequence similarity (Makarova *et al*., 2020), but structural analyses revealed that they have similar structures and bind to similar positions in the Cas protein complexes (Hayes *et al*, 2016; Hu *et al*, 2022; Jackson *et al*, 2014; Jore *et al*., 2011; Lu *et al*., 2024; Mulepati *et al*, 2014; Wiedenheft *et al*., 2011). Structural studies for the type I-E Cascade complex have shown that Cas11 is present in two copies as “belly” located opposite the Cas7 multimers (Fig. 1A.) (Hayes *et al*., 2016; Jackson *et al*., 2014; Jore *et al*., 2011; Mulepati *et al*., 2014; Wiedenheft *et al*., 2011; Zhao *et al*, 2014). Structural analyses of the type I-A Cascade have revealed that five Cas11 subunits form the inner belly (Hu *et al*., 2022). However, data reported so far about the different Cas11 proteins suggest that despite their structural similarity they do have differing functional roles in the interference (Liu & Doudna, 2020; Reeks *et al*, 2013; Tan *et al*, 2022).

**Figure 1.**
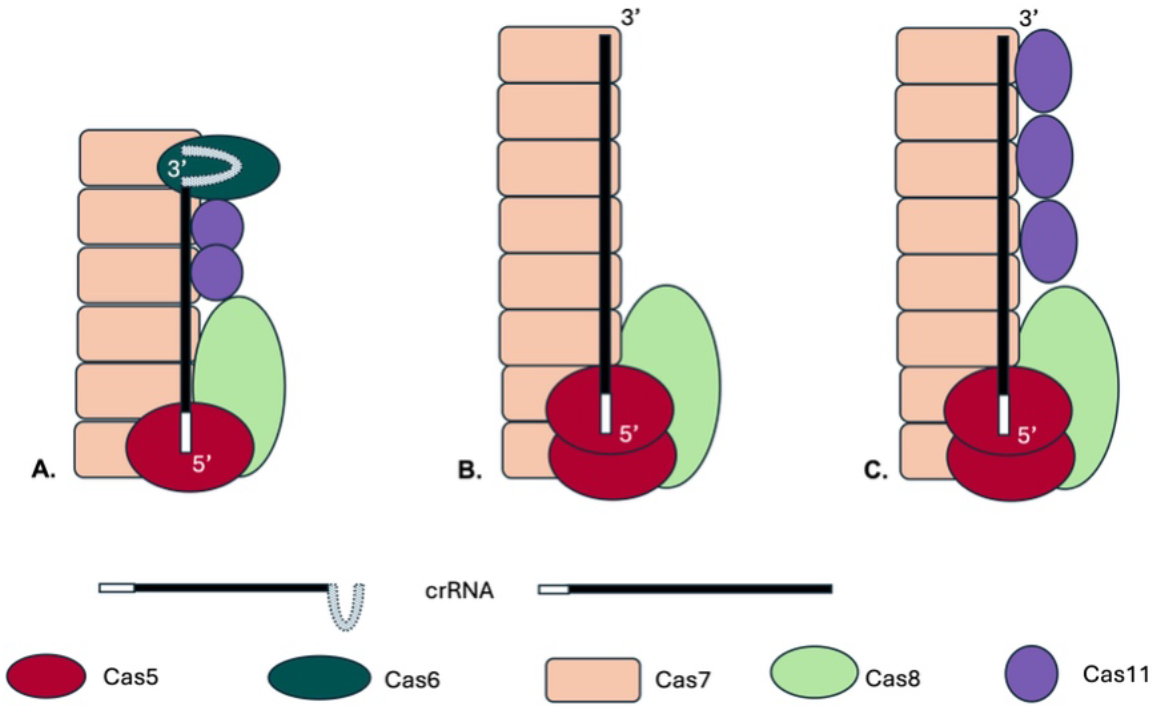
Schematic drawings of Cascade complexes. The type I CRISPR-Cas complexes contain a large subunit (Cas8 or Cas10), Cas7 proteins form the complex’s backbone and Cas5 binds at the crRNA 5’ end. With the exception of type I-F all Cascades contain as small subunit, the Cas11 protein. **A**. The type I-E Cascade complex contains 6 Cas7 proteins, one Cas5, Cas6 and Cas8 each and two Cas11 proteins (in purple) (Jore *et al*, 2011; Wiedenheft *et al*, 2011). **B**. A potential complex structure of the *Haloferax* Cascade complex is shown. The structure of the *Haloferax* Cascade complex has not yet been solved, but the composition of a purified Cascade was determined (Brendel *et al*, 2014). In this previous work potential Cas11b and Cas8b peptides could not be distinguished therefore no Cas11b is shown. **C**. A potential *Haloferax* Cascade structure including the here identified Cas11b is shown, the number of Cas11b proteins was assumed as found in the bacterial type I-B (Lu *et al*, 2024).

Recently, it has been discovered during analysis of the bacterial type I-D system that the gene for the large protein (Cas10) contains an internal translation initiation site near the 3’ end, resulting in an internal in-frame translation of a Cas11 protein (McBride *et al*, 2020). Further analyses showed that similar internal translation initiation sites also exist in the genes for the large Cas proteins (Cas8) of the bacterial type I-B and I-C systems (Hussain *et al*, 2023; McBride *et al*., 2020; O’Brien *et al*, 2020; Tan *et al*., 2022). Since all reports so far concentrated on bacterial systems and the archaeal *Haloferax* type I-B CRISPR-Cas system does not contain a separate gene for a Cas11b protein, we investigated here whether the *cas8b* gene encodes a Cas11b protein. And if so how Cas11b contributes to Cascade stability and function and if it has additional cellular functions beyond virus defence.

## Materials and Methods

### Strains and growth conditions

Strains, plasmids and primers used are listed in Suppl. Table 1. *Hfx. volcanii* H119 was used as parent strain (wild type strain) for all experiments. H119 is derived from wild type strain DS2 by removal of plasmid pHV2 and deletion of three genes that can be used as selection markers (Δ*pyrE2*, Δ*trpA*, Δ*leuB*) (Allers *et al*, 2004). All *Hfx. volcanii* strains were grown aerobically with shaking (200 rpm) at 45 °C in Hv-YPC (rich medium) or Hv-Cab media (minimal medium) with appropriate supplements if carrying any plasmid as described (Allers *et al*., 2004).

*E. coli* strains DH5α (Invitrogen, Thermo Fischer Scientific, Waltham, MA, USA) and GM121 (Allers *et al*, 2010) were grown aerobically at 37°C in 2YT medium (Miller, 1972) supplemented with 75 µg/ml ampicillin at 37 °C for selection.

### Cloning of plasmids

The generation of plasmid pTA927-*cas8b*-FLAGN for expression of the Cas8b protein under control of the p.*tna* promotor carrying an N-terminal 3xFLAG-tag is described in (Cass *et al*, 2015). Plasmid pTA927-*cas8b*-FLAGC for expression of the Cas8b protein under control of the p.*tna* promotor carrying a C-terminal 3xFLAG-tag, was generated by amplification of the *cas8b* gene using the oligonucleotides 5-Csh1-NdeI and 3-Cas8-SmaI and pTA927-*cas8b*-FLAGN as template, the resulting DNA fragment was digested with *Nde*I and *Sma*I and ligated into *Nde*I and *SnaB*I digested vector pTA927-FLAGC (Hadjeras *et al*, 2023), resulting in the final construct pTA927-*cas8b*-FLAGC. The plasmid for expressing FLAG-Cas8bM545A was generated by performing a mutagenesis PCR using the QuikChange II XL Site-Directed Mutagenesis Kit (Agilent Technologies) with the oligonucleotides 8M545A and 8M545Ar and pTA927-*cas8b*-FLAGC as template, resulting in the final construct pTA927-*cas8b*M545A-FLAGC. Plasmid pTA927-cas11b for expression of the Cas11b protein under the control of the p.*tna* promoter was generated as follows. The *cas11b* gene was amplified by PCR using oligonucleotides 5-Cas11-NdeI and 3-Cas11-XbaI and pTA927-*cas8b*-FlagC as template. The resulting PCR product was cleaved with *Nde*I and *Xba*I and cloned into vector pTA927 (also cleaved with *Nde*I and *Xba*I) (Allers *et al*., 2010), resulting in the final construct pTA927-*cas11b*. To obtain the plasmid for expression of the mutated Cas8b protein (lacking the start codon for Cas11b) and the Cas11b from the same plasmid, the *cas11b* gene under the control of the p.*tna* promoter was amplified by PCR using the oligonucleotides 5-ptna-BamHI and 3-cas11-XbaI and pTA927-*cas11b* as template and cloned into the plasmid pTA927-*cas8bM545A*-FlagC via the restriction sites *BamH*I and *Xba*I, resulting in the final construct pTA927-*cas8bM545A*-FlagC-*cas11b*. The plasmid for generating the *cas8bM545A* mutant strain was generated as follows. The mutated *cas8bM545A* gene was amplified by PCR using the oligonucleotides 5-cas8-fw and 3-cas8 rev and pTA927-*cas8bM545A*-FlagC as template. The existing plasmid pTA131-up.do(Csh1) (Cass *et al*., 2015), which contains the app. 500-550 basepairs upstream and downstream regions of the *cas8b* gene, was used as template in a PCR reaction using the oligonucleotides IP cas8 rev and IP cas8 fw to amplify the plasmid in an inverted PCR. The mutated cas8bM545A insert was then ligated into that vector, resulting in the final construct pTA131-up.*cas8b545A*.do.

### Generation of the cas8bM545A strain

To generate the *cas8bM545A* mutant strain a Δ*cas8b* deletion strain was transformed with plasmid pTA131-up.Cas8bM544A.do to integrate the modified gene into the *cas8b* gene locus using the pop-in/pop-out method (Allers *et al*., 2004; Bitan-Banin *et al*, 2003). Pop-out-candidates were validated by Southern blot analysis as follows (Suppl. Fig. 1). 10 µg of isolated genomic DNA were digested with *Sac*II and resulting fragments were separated by size on an 0.8 % agarose gel and transferred to a nylon membrane Hybond^™^-N+ (GE Healthcare) via capillary blot. For detection of DNA fragments, two PCR generated probes were used: one for detection of the *cas8b* gene (Cas8_intern) and one for the downstream region (Cas8_do). Probes were labeled with [γ-^32^P]-dCTP using the DECAprime II DNA labeling kit (Thermo Fisher Scientific) and used as hybridisation probe.

### Identification of Cas11b peptides by mass spectrometry

SDS-PAGE-separated protein samples were processed as described by (Shevchenko *et al*, 1996). The resulting peptides were loaded onto nano-HPLC (Dionex Ultimate 3000 UHPLC Thermo Fischer Scientific) coupled with in-house packed C18 column (ReproSil-Pur 120 C18-AQ, 3 µm particle size, 75 µm inner diameter, 30 cm length, Dr. Maisch GmbH). The peptides were separated with a linear gradient of 11– 42% buffer B (80% acetonitrile and 0.1% formic acid) at flow rate of 300 nL/min over 43 min gradient time. Eluting peptides were analysed by Q Exactive mass spectrometer (Thermo Fischer Scientific). The MS data was acquired by scanning the precursors in mass range from 350 to 1600 m/z at a resolution of 70,000 at 200 m/z. Top 20 precursor ions were chosen for MS2 by using data-dependent acquisition (DDA) mode at a resolution of 17,500 at 200 m/z. Data analysis and search was performed using Mascot Software (2-3-02) and Scaffold 5.2.1 with Uniprot_Haloferax_volcanii database with 11678 entries.

Peptide identification and quantification was done with MaxQuant version 2.2.0.0 (Cox and Mann, 2008) with default settings against *Haloferax volcanii* database accessed from UniProt at 24.01.2023 (UP000008243; 3921 entries) and the sequence of the predicted Cas11b protein. The resulting peptides.txt table was used to extract raw intensity values for the selected peptides.

### Interference test

The invader plasmid pTA352-PAM3-P1.1 was used to test the interference activity of the endogenous CRISPR-Cas type I-B system (Brendel *et al*., 2014; Fischer *et al*., 2012). This plasmid is based on the *Haloferax* shuttle vector pTA352 (Norais *et al*, 2007) including spacer 1 of the CRISPR locus P1 and the PAM sequence TTC (PAM3) (Fischer *et al*., 2012). As a control reaction *Haloferax* cells were transformed with the vector pTA352 without insert. Plasmids were passaged through *E. coli* GM121 cells (to avoid methylation) and were then introduced into *Haloferax* cells using the PEG method (Allers *et al*., 2004; Cline *et al*, 1989). Four dilutions of the transformed cells (undiluted, dilutions 1:10, 1:100 and 1:1000) were spotted on plates with suitable selection medium (minimal medium containing leucin). To confirm active interference, *H. volcanii* cells were transformed at least three times with the plasmid invader or the control vector.

### Northern blot hybridisation

To analyse crRNA levels, H119 x pTA927, Cas8bM545A x pTA927 and Cas8bM545A x pTA927-Cas11 cells were grown in Hv-Cab supplemented with uracil until OD_650_ of 0.49-0.54. Total RNA was isolated using NucleoZol (Macherey-Nagel) and 10 µg of each culture were separated on an 8% denaturating polyacrylamide gel. RNA was transferred to a nylon membrane Hybond^™^-N+ (GE Healthcare) by electroblotting and the membrane was hybridised overnight at 50 °C with oligo P1SP1, which was labeled with γ-ATP-^32^P using T4 polynucleotide kinase (Thermo Scientific). The intensity of the radioactive signal was recorded using an Imaging Plate BAS-MS (Fujifilm) and a Amersham Typhoon-imager (Cytivia). The same procedure was repeated with an oligo binding to the 5S rRNA as a loading control to normalise the P1SP1 signal. The P1SP1 probe detects spacer1 of the CRISPR locus P1.

### Size exclusion chromatography

Wild-type strain H119 and mutant strain *cas8b*M545A were transformed with pTA927-cas7-N-Flag. As a control H119 was transformed with pTA927-FLAGcontrol. Cells were grown in 500 ml Hv-Cab medium to OD_650_ of 0.78-0.86, then tryptophane was added to a final concentration of 3 M to induce expression of Cas7-N-Flag. Cells were grown for 3 hours, harvested and subsequently washed in enriched PBS (2.5 M NaCl, 150 mM MgCl_2_, 1x PBS). After lysis by ultrasonification in 10 ml Lysis Buffer (100 mM Tris-HCl pH 7.5, 10 mM EDTA, 150 mM NaCl), membranes and unsoluble material were pelleted by centrifugation at 100,000 xg at 4°C for 1 hour. The Flag-tagged proteins and their interaction partners were further purified from the supernatant with Anti-Flag M2 Affinity Gel (Sigma Aldrich), protein concentration was determined with Roti-Nanoquant (Carl Roth GmbH). 200 µg/sample or the complete fraction were filtered using an Amicon Ultra Centrifugation filter 50 kDa MWCO (Merck). The material retained on the filter was resuspended in 300 µl SEC-Buffer (1 M NaCl, 25 mM Tris-HCl pH=7.0, 5 mM MgCl_2_), and then loaded on a SuperdexTM 200 Increase 10/300 GL column (Cytivia) with a flow-rate of 0.375 ml/min. The flow through was collected in 1 ml fractions and conductivity as well as absorption at 280 nm were measured. Proteins of the first fractions were separated on a 10 % SDS-Gel and transferred to a nitrocellulose membrane (Roti®NC, Carl Roth GmbH) by semi-dry western blot. The FLAG-tag containing proteins were visualised using an Anti-FLAG M2-mcAb-HRP antibody (Sigma Aldrich).

### RNAseq

Five biological replicates of the wildtype strain H119 and the mutant strain *cas8b*M545A were grown in YPC at 45 °C to an OD_650_ of 0.48 - 0.53. Total RNA was isolated using NucleoZol™(Macherey-Nagel, Düren, Germany). A Turbo-DNase digest (Thermo Fisher Scientific) was performed followed by depletion of ribosomal RNA molecules using a commercial Pan-Archaea riboPOOL rRNA depletion kit (siTOOLs Biotech, dp-K024-000027). The ribo-depleted RNA samples were first fragmented using ultrasound (2 pulses of 30 s at 4°C). Then, an oligonucleotide adapter was ligated to the 3’ end of the RNA molecules. First-strand cDNA synthesis was performed using M-MLV reverse transcriptase with the 3’ adapter as primer. After purification, the 5’ Illumina TruSeq sequencing adapter was ligated to the 3’ end of the antisense cDNA. The resulting cDNA was PCR-amplified using a high-fidelity DNA polymerase and the barcoded TruSeq-libraries were pooled in approximately equimolar amounts. Sequencing of pooled libraries, spiked with PhiX control library, was performed at 7-14 million reads per sample in single-ended mode with 100 cycles on the NextSeq 2000 platform (Illumina). Demultiplexed FASTQ files were generated with bcl-convert v4.3.6 (Illumina). Raw sequencing reads were subjected to quality and adapter trimming via Cutadapt (Martin, 2011) v2.5 using a cutoff Phred score of 20 and discarding reads without any remaining bases (parameters: --nextseq-trim=20 -m 1 -a AGATCGGAAGAGCACACGTCTGAACTCCAGTCAC). Afterwards, all reads longer than 11 nt were aligned to the *Haloferax volcanii* DS2 reference genome (RefSeq assembly accession: GCF_000025685.1 excluding plasmid pHV2 (NC_013965.1)) using the pipeline READemption (Förstner *et al*, 2014) v2.0.3 with segemehl v0.3.4 (Hoffmann *et al*, 2009) and an accuracy cut-off of 95% (parameters: -l 12 -a 95). READemption gene_quanti was applied to quantify aligned reads overlapping genomic features by at least 10 nt (-o 10) on the sense strand based on annotations (CDS, mature_transcript, rRNA, tRNA) originating from a detailed continuous manual curation effort (Laass *et al*, 2019). The annotation version from 28-MAR-2023 was used (annotated protein sequences in: https://doi.org/10.5281/zenodo.7794769), this version includes small ORFs reported in (Hadjeras *et al*., 2023). Based on these counts, differential expression analysis was conducted via DESeq2 (Love *et al*, 2014) v1.24.0. Read counts were normalized by DESeq2 and fold-change shrinkage was conducted by setting the parameter betaPrior to TRUE. Differential expression was assumed at adjusted p-value after Benjamini-Hochberg correction (padj) < 0.05 and |log2FoldChange| ≥ 2 (Table 1).

**Table 1.**
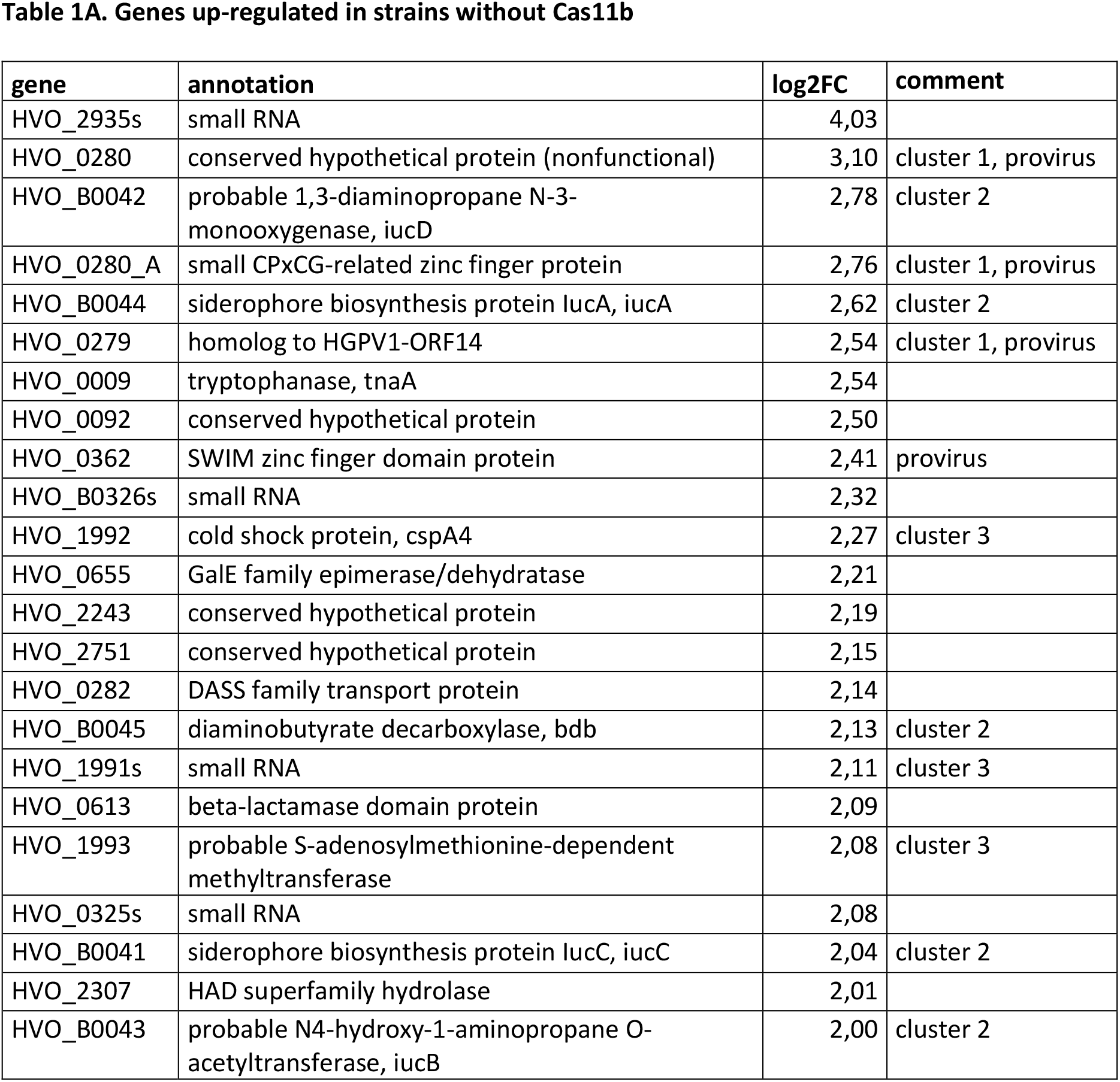
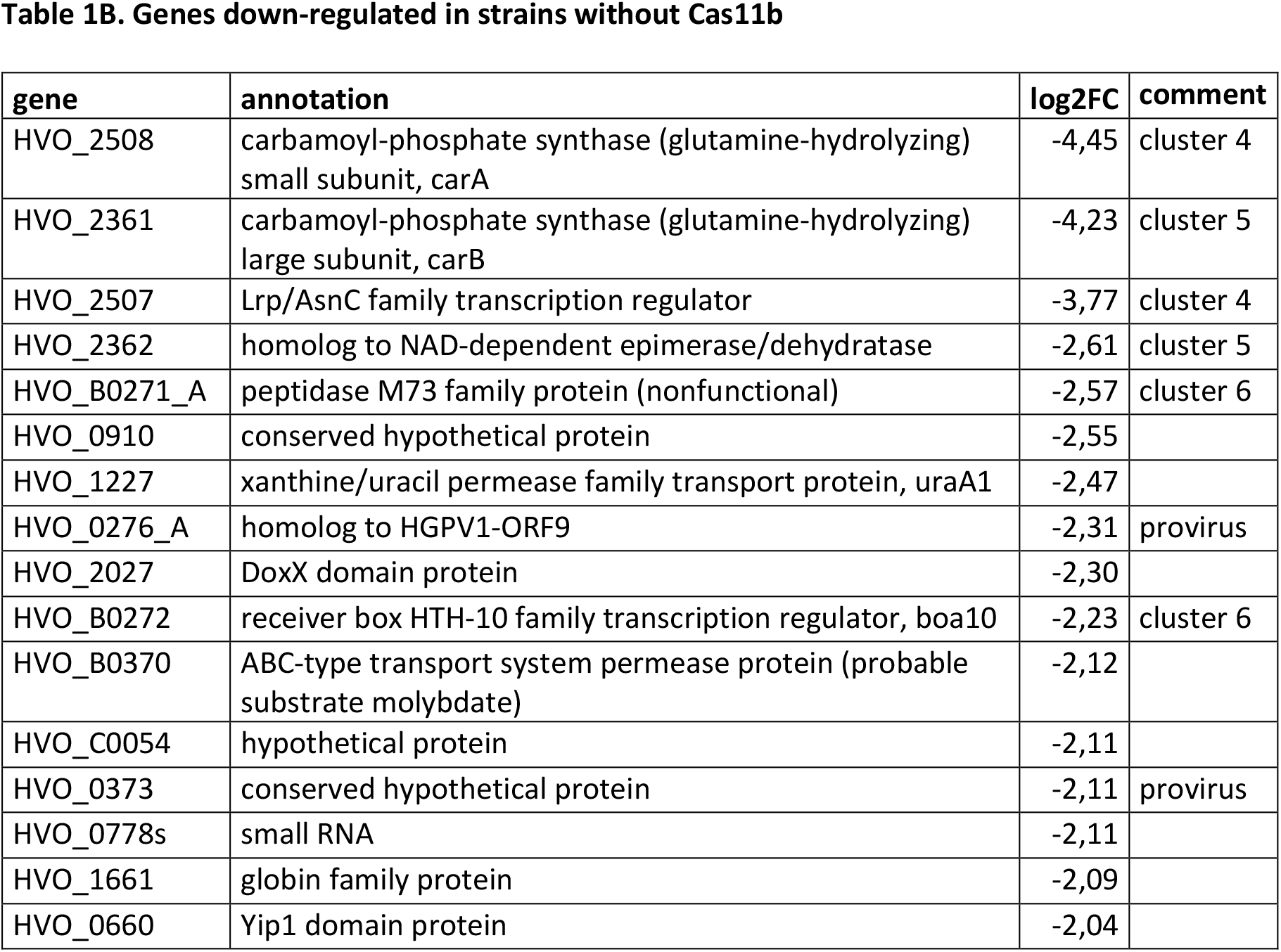
Genes up-regulated in strains without Cas11b. Transcriptomes of the wild type *Haloferax* strain (H119) and the *cas8*M545A mutant have been determined and compared. **A**. Genes up-regulated with a log2 fold change of more than 2.0 are shown. **B**. Genes down-regulated with a log2 fold change of less than -2.0 are shown. Only genes that show significant changes (padj < 0.05) are shown. Regulation of pHV4 genes, most of them down-regulated, is shown in Suppl. Table 2. Genes that are part of one of the provirus regions and neighbouring genes are indicated (column comment).

## Results

### Cas11b is encoded in frame by the *cas8b* gene

As initial approach to determine expression of a Cas11b protein from the *cas8b* gene we generated a plasmid, which expresses a Cas8b-FLAG fusion protein. Upon expression of an N-FLAG-Cas8b fusion protein only a single band was visible on the western blot (Fig. 2A. and C.). In contrast, two signals were detected upon expression of a C-FLAG-Cas8b fusion protein (Fig. 2B. and C.), suggesting that a second protein of about 29 kDa is produced with a translation start in the 3’ part of the gene. Based on the sizes observed, the long protein probably corresponds to Cas8b (calculated molecular weight 92.8 kDa) while the smaller protein is likely to be Cas11b translated from the *cas8b* open reading frame starting at the ATG for Met545 (calculated molecular weight 28.9 kDa) (Fig. 3).

**Figure 2.**
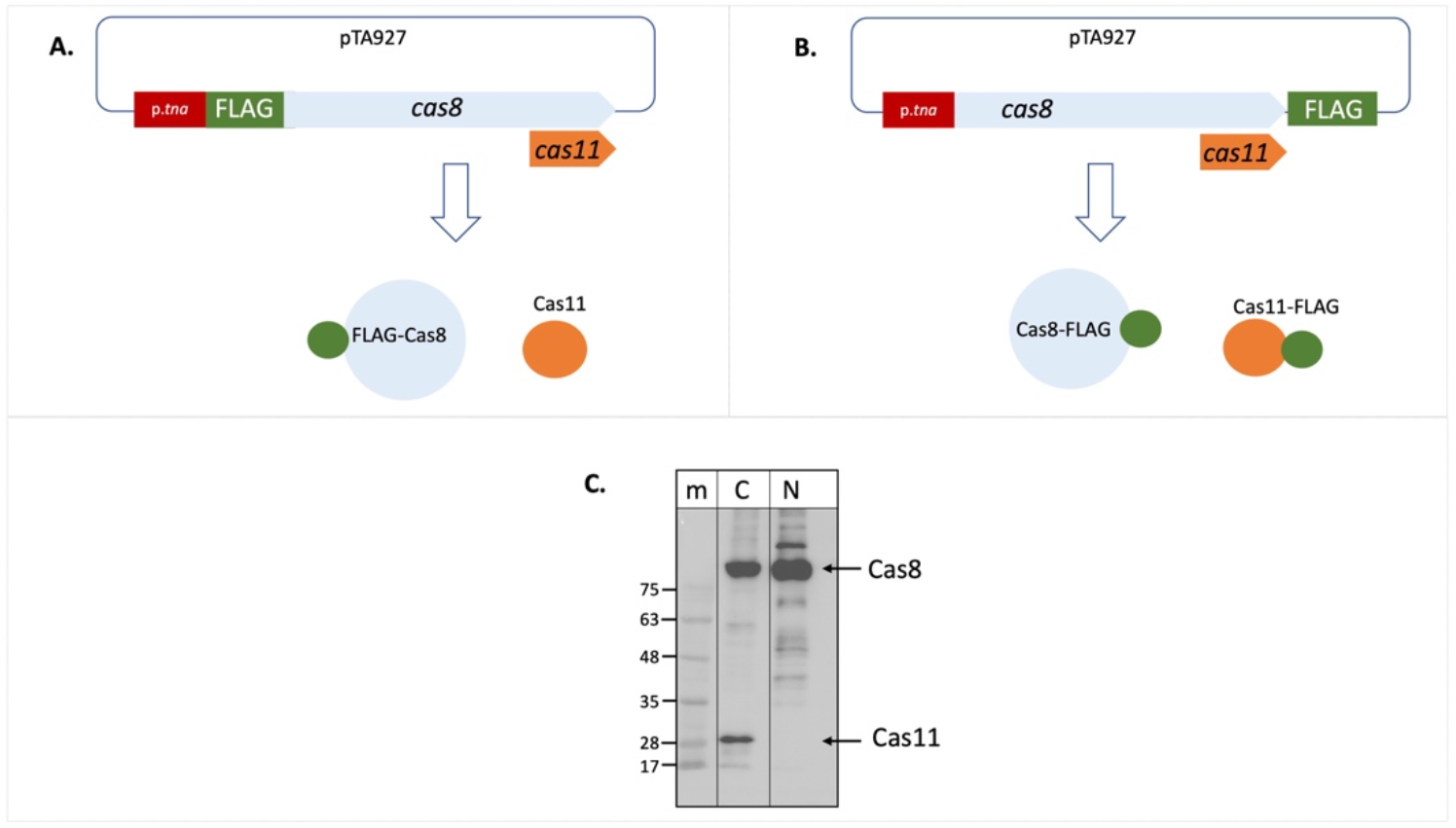
Cas11b is encoded in the *cas8b* gene. The cDNA for the FLAG tag was fused to the *cas8b* gene to allow expression of a Cas8b-FLAG protein. **A**. Fusion to the N-terminus results in a FLAG-Cas8b protein and an untagged Cas11b protein (not visible in the western). **B**. Fusion to the C-terminus results in a Cas8b-FLAG protein as well as a Cas11b-FLAG protein. **C**. *Haloferax* wild type cells (H119) were transformed with both constructs. Proteins were isolated and separated on SDS PAGE and subsequently transferred to a western membrane that was probed with an Anti-FLAG antibody. Lane m: protein size marker, sizes are given at the left in kDa, lane N: FLAG fused to the N-terminus of the protein, lane C: FLAG fused to the C-terminus of the protein. The calculated molecular weight of Cas8b is 79.4 kDa and of Cas11b (starting at Met545) is 25.5 kDa. Due to the high amount of acidic amino acids, halophilic proteins run generally slower on SDS PAGE (Guan *et al*, 2015). Taking this into account, the calculated molecular weight of Cas8b is 92.8 kDa and of Cas11b (starting at Met545) is 28.9 kDa, their position in the gel is indicated with arrows at the right.

**Figure 3.**
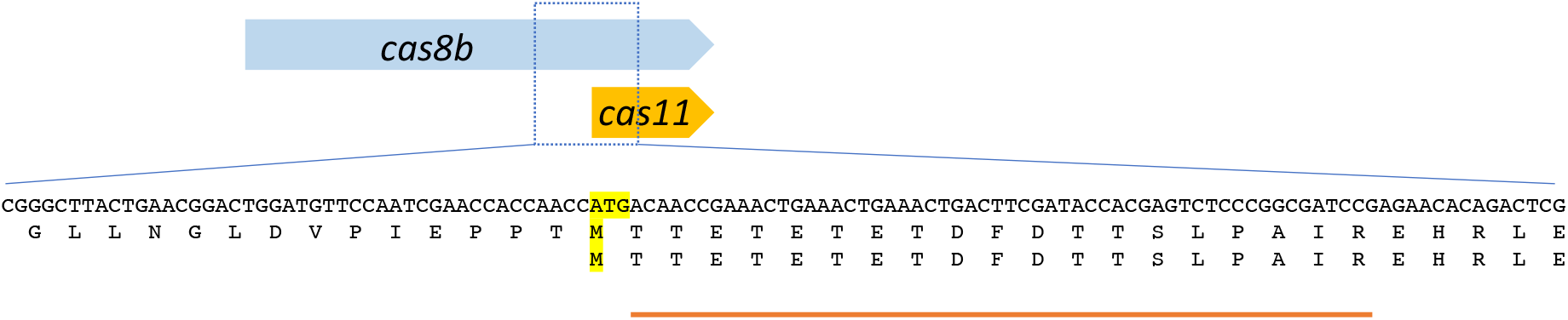
The *cas8b* gene encodes Cas8b and Cas11b. The Cas11b protein is encoded in the 3’ terminal third of the *cas8b* gene. The translation start ATG and the corresponding methionine residues (the internal Met545 of the Cas8b sequence (upper sequence) and the start codon of Cas11b (lower sequence)) are highlighted in yellow. The amino acid sequence of Cas8b and Cas11b is shown. The peptide identified in the mass spectrometry analysis is indicated by an orange line. The observed peptide starts with Thr546, consistent with the ATG for Met545 being used for translation initiation with subsequent cleavage of the N-terminal methionine, which is a common in step haloarchaeal N-terminal protein maturation.

Both proteins, the small one with 29 kDa and the large one with 93 kDa, were analysed by mass spectrometry. The larger one was confirmed to be Cas8b and there is a strong indication, that the smaller protein is Cas11b (Fig. 3, Suppl. Fig. 2). A peptide starting at Thr546 was identified, strongly indicating that translation initiated at the ATG for Met545 with subsequent cleavage of the N-terminal Met, which is very often found in haloarchaea (Falb *et al*, 2006; Schulze *et al*, 2020). This peptide cannot be generated from Cas8b by tryptic cleavage, however it could be a result of unspecific fragmentation. Therefore, we wanted to obtain further support for a Cas11b translation initiation at Met545. To further test the hypothesis of an internal translation start and to prove that Met545 is the start codon of Cas11b, we mutated the Met545 codon (ATG) to Ala545 (GCG).

The mutated *cas8bM545A* gene was cloned as C-FLAG-fusion construct (Fig. 4). Expression of the C-FLAG-cas8bM545A yielded only a single signal corresponding to a protein of about 93 kDa in the western. Thus, the Met545Ala mutation prevents translation initiation for the Cas11b protein and therefore 29 kDa protein can not be detected in the western. Taken together these data show that a Cas11b protein is translated from the *cas8b* gene with an internal in-frame translation start.

**Figure 4.**
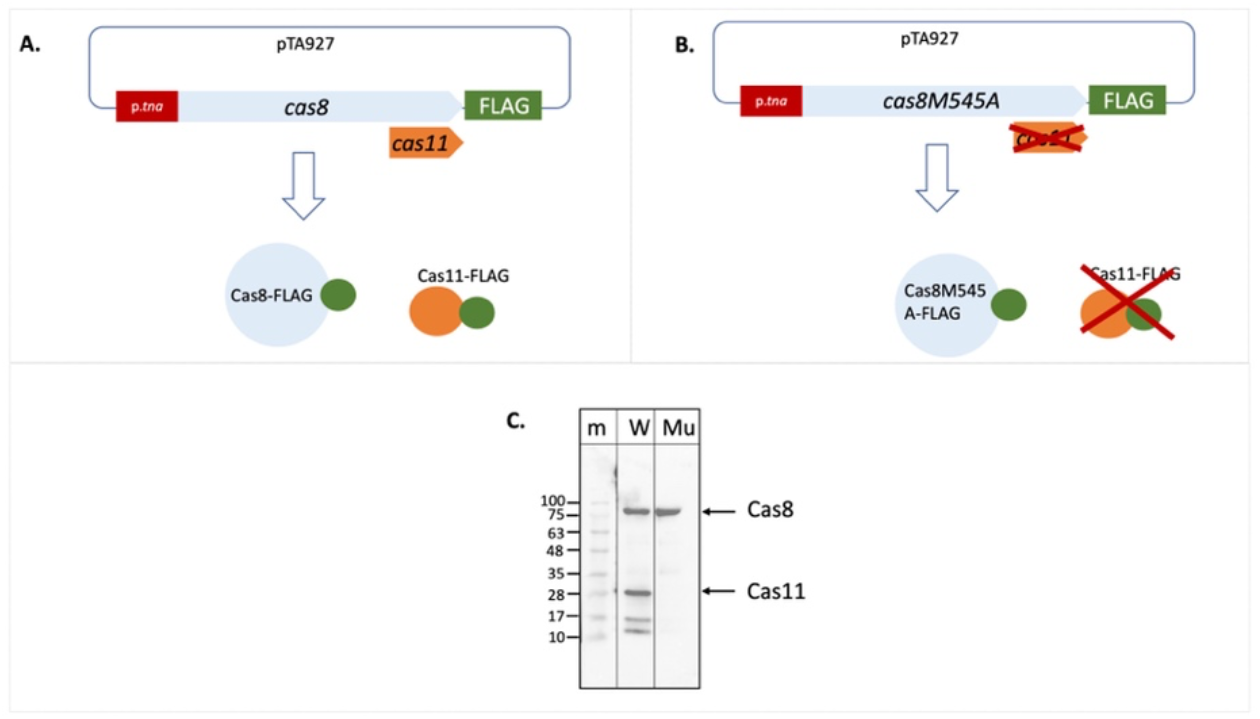
The Cas8bM545A mutation abolishes production of Cas11b. The FLAG cDNA was cloned in frame downstream of the wild type *cas8b* gene and also of the mutated *cas8b*M545 gene (**A**. and **B**.). Expression of this gene results in a Cas8b-FLAG fusion protein and potentially also in a Cas11b-FLAG fusion protein. **C**. *Haloferax* cells were transformed with the plasmids, proteins were extracted and separated on a SDS PAGE. After transfer of the proteins to a western membrane the FLAG tag was identified with Anti-FLAG antibodies. Lane m: protein size marker, sizes are given at the left in kDa, lane W: C-FLAG-Cas8b and C-FLAG-Cas11b are expressed from the wild type *cas8b* gene, lane Mu: Only C-FLAG-Cas8b is expressed from the mutated *cas8b* gene. The calculated molecular weight of Cas8b is 92.8 kDa and of Cas11b (starting at Met545) is 28.9 kDa (taking into account the high percentage of acidic amino acids), their position in the gel is indicated with arrows at the right.

### Cas11b is required for an efficient interference reaction

Next we wanted to examine whether Cas11b is required for the defence activity of the CRISPR-Cas type I-B system. An effective interference test using an invader plasmid was previously established for the *Haloferax* type I-B system (Fischer *et al*., 2012). A typical invader plasmid contains a protospacer flanked by a protospacer adjacent motif (PAM) that is detected by one of the crRNAs (Maier *et al*., 2019; Turgeman-Grott *et al*, 2019). Here, we used an invader plasmid containing the functional PAM TTC and protospacer P1-1 that is detected by the *Haloferax* CRISPR-Cas system, since the first spacer from CRISPR locus P1 can basepair to it (Fischer *et al*., 2012). To confirm that expression of a Cas8b-FLAG fusion protein from a plasmid is functionally active in the Cascade complex we transformed a Δ*cas8b* strain with a plasmid expressing the Cas8b-C-FLAG fusion protein and subsequently with the invader plasmid. The invader tests clearly showed that the CRISPR-Cas system is active in destroying the invader plasmid (Fig. 5A.).

**Figure 5.**
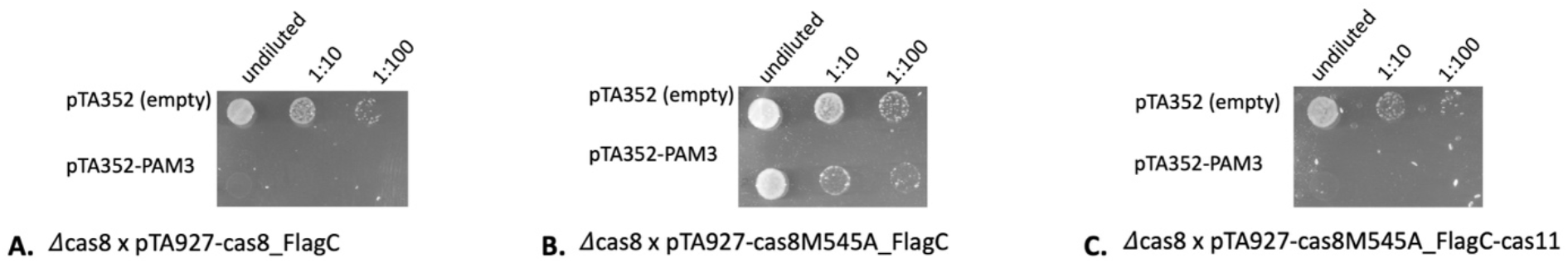
Without Cas11b the interference reaction is less efficient. *Haloferax* cells were transformed with an invader plasmid (pTA352-PAM3-P1.1) and plated on selective medium. Only cells that did not degrade the invader plasmid can grow. As control cells are transformed with the pTA352 plasmid. **A**. Cells with Cas8b and Cas11b (Δcas8 x pTA927-cas8_FlagC) efficiently degrade the plasmid invader. **B**. Cells with Cas8b but without Cas11b (Δcas8 x pTA927-cas8M545A_FlagC) cannot degrade the plasmid invader anymore. **C**. When the mutant Cas8b and additionally Cas11b (Δcas8 x pTA927-cas8M545A_FlagC_FlagC-cas11) are present, cells efficiently degrade the plasmid invader.

In contrast, when the deletion strain Δ*cas8b* was transformed with a plasmid expressing the Cas8bM545A mutant the interference reaction was not active anymore, and there was no reduction of transformation efficiency evident compared to the control (Fig. 5B.).

Thus, without Cas11b the interference reaction is far less efficient. Upon transformation of Δ*cas8b* with a plasmid expressing both, mutated Cas8b as well as Cas11b independently (Δ*cas8b* x pTA927-cas8M545A_FLAGC-cas11), interference activity was recovered, and a reduction of transformation efficiency was again observed (Fig. 5C.). Thus Cas11b is required for an efficient defence reaction.

### Cas11b and the stability of crRNAs

To investigate further why Cas11b is important for efficient interference, we investigated the crRNA levels in a Cas11b less strain. The crRNA is one of the key players of the CRISPR-Cas system and important for guiding the Cascade complex to the target. Previous studies showed that crRNAs are not stably maintained in *H. volcanii* cells when Cascade is not present, suggesting that these small RNAs are protected by binding to the Cas protein complex (Brendel *et al*., 2014). To analyse whether Cas11b is required for stable crRNA maintenance we investigated the presence and levels of crRNAs in a Cas11b-deficient strain using northern blot analyses (Suppl. Fig. 3). Wild type *Haloferax* cells transformed with plasmid pTA927 (without an insert) express the native Cas8b and Cas11b from the original chromosomal *cas8b* gene. In these cells crRNAs are readily detected. *Haloferax* mutant cells (*cas8*M545A), which express only the mutated Cas8b contain similar levels of crRNA. And complementation of the *Haloferax* mutant cells (*cas8*M545A) with pTA927-*cas11b* also shows wild type levels of crRNAs. These data show that crRNAs are not affected when Cas11b is missing.

### Slightly more Cascade complexes can form in the presence of Cas11b

Since Cas11b was shown to be part of Cascade in other systems we wanted to investigate whether Cascade without Cas11b is still stable. *Haloferax* cells with Cas11b (Δcas7xpTACas7FLAG) and without Cas11b (*cas8*M545AxpTACas7-FLAG) were grown, the soluble protein fraction was isolated, and proteins were separated on a gel filtration column after a FLAG-purification step. Comparison of the complex levels between the two strains showed that in the absence of Cas11b slightly less Cascade complexes were formed and eluted similarly to previous analyses (Brendel, 2014) (Fig. 6, Suppl. Fig. 4). Analysis of the fractions with western blot confirmed the presence of Cas7-FLAG (Suppl. Fig. 4).

**Figure 6.**
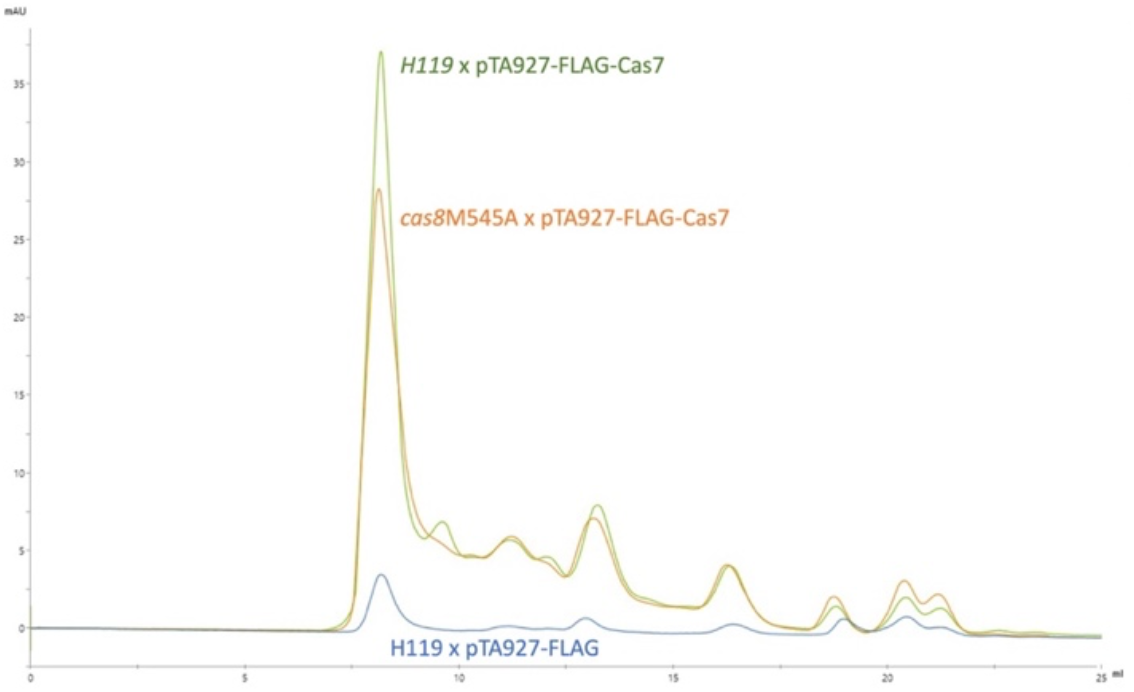
Cascade with Cas11b is slightly more stable. Cascade complexes from *Hfx. volcanii* can be purified with gel filtration as reported previously (Brendel, 2014). If Cas11b is present the amount of Cascade complexes is higher (green curve) than without (orange curve). Blue curve: only the FLAG peptide was expressed, purified and loaded onto the gel filtration column. Green curve: (Δcas7x pTA927-Cas7FLAG) cells were grown, FLAG-Cas7 and interacting proteins were purified with FLAG agarose and loaded onto the gel filtration column. Orange curve: FLAG-Cas7 was purified from *cas8*M545A x pTA927-Cas7-FLAG cells. The elution volume is shown in ml on the x-axis and the absorption is shown in milli absorption units (mAU) on the y-axis.

### Without Cas11b a plethora of changes in gene expression is observed

We next wanted to determine how the elimination of Cas11b affects the transcriptome of the cell. RNA was isolated from wild type (expressing Cas8b and Cas11b) and *cas8*M545A cells (expressing only the mutated Cas8bM545A but not Cas11b) and used for RNA-seq analyses. Many genes showed a change in expression (Table 1). Genes encoded on the mini-chromosome pHV4 were especially affected (Suppl. Table 2): more than 100 genes between HVO_A0279_A and HVO_A0405 were down-regulated, only six pHV4 genes were up-regulated. Genes from the encoded proviruses Halfvol1 and Halfvol2 were also regulated (down-as well as up-regulated). The expression of five small RNAs as well as of 13 genes coding for proteins without known function also changed significantly.

The five neighbouring genes HVO_B0041-45 (*iuc*C,D,B,A and bdb) were up-regulated with a log2fold more than 2; these genes are believed to be involved in the siderophore biosynthetic pathway, which is important for iron acquisition in iron-poor environments (Blin *et al*, 2023; Sailer *et al*, 2024). Removal of Cas11b thus has an effect on this iron related pathway, probably relieving repression during growth in iron-replete medium.

## Discussion

### One gene two proteins

In *Haloferax* Cas11b is made from an internal in-frame translation initiation site present in the *cas8b* gene. To our knowledge this is the first experimental proof for an internal in-frame translation in archaea. A ribosome profiling study performed with *H. volcanii* reported potential internal in frame translation initiation sites (Gelsinger *et al*, 2020), but the Cas11b start site was not detected in this study and actual proof for an internal translation initiation was not shown. This is also the first report about an archaeal type I-B encoding Cas11b within the gene for the large subunit Cas8b. A short fragment of Cas8b was previously identified for the type I-B systems of *Clostridium thermocellum* und *Methanococcus maripaludis*, but here fragmentation of the Cas8b protein was suggested (Richter *et al*, 2017). Structural studies of other type I Cascade complexes showed that they contain varying numbers of Cas11 proteins: five (I-A), three (I-B) or two (I-C, I-D, I-E (Hu *et al*., 2022; Jackson *et al*., 2014; Jore *et al*., 2011; Lu *et al*., 2024; McBride *et al*., 2020; Mulepati *et al*., 2014; O’Brien *et al*., 2020; Schwartz *et al*, 2022; Wiedenheft *et al*., 2011; Zhao *et al*., 2014). This suggests that for the I-B and I-D systems translation from the internal AUG must be three times and twice, respectively, as effective as from the Cas8b AUG.

Unfortunately, the presence of a Shine-Dalgarno sequence can not be used as an additional proof for an internal translation start, since in contrast to bacterial mRNAs, haloarchaeal mRNAs do not always contain Shine Dalgarno sequences and especially in *H. volcanii* the number of mRNAs containing a Shine-Dalgarno sequence is very low (Kramer *et al*, 2014). Thus, a missing Shine-Dalgarno sequence does not argue against the presence of a translation start. However, a potential SD sequence (GGATG instead of the standard GGAGG (Kramer *et al*., 2014)) can be identified upstream of the Cas11b M545 (Fig. 8).

**Figure 8.**
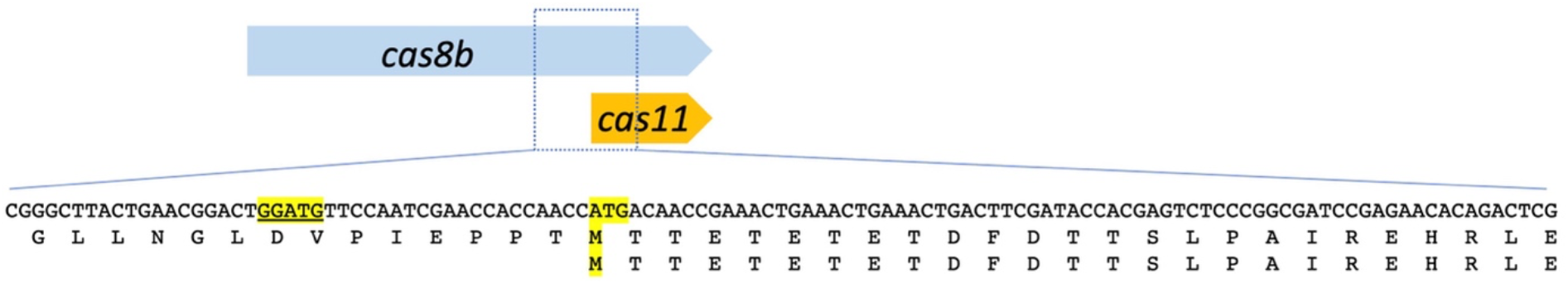
A potential Shine-Dalgarno sequence upstream of the Cas11b ATG. The Cas11b protein is encoded in the 3’ terminal third of the *cas8b* gene. The translation start ATG and the corresponding methionine residues (M545) are highlighted in yellow. The amino acid sequence of Cas8b and Cas11b is shown below the DNA sequence. A potential Shine Dalgarno sequence is located upstream of the M545 ATG: GGATG (highlighted in yellow and underlined; the standard SD for Hfx. volcanii is: GGAGG (Kramer *et al*., 2014)).

### Complex formation without Cas11b

Our data show that Cascade can form in *Haloferax* without Cas11b, which is in line with the data reported for the I-D Cascade complex (McBride *et al*., 2020) and for the *Synechocystis* sp. I-B Cascade (Tan *et al*., 2022). The amount of Cascade complexes without Cas11b reported here seems to be slightly lower than with Cas11b but complexes are still able to assemble. In contrast type I-C Cascade does not form without Cas11c (Tan *et al*., 2022). Thus, for the *Haloferax* type I-B Cascade stability Cas11b is not essential. Recently an anti-CRISPR protein was found, that chips away the Cas7 subunits of a type I-F Cascade complex (Trost *et al*, 2024). Cascades from type I-F do not have Cas11, which probably makes the observed removal of Cas7 proteins easier for the anti-CRISPR protein.

### Cas11b and interference efficiency

The interference reaction is drastically reduced without Cas11b. Although Cascade complexes still form without Cas11b, they are not active in interference anymore. A loss in interference was also observed for the *Leptospira interrogans* I-B system upon depletion of the Cas11b (Hussain *et al*., 2023). Similar observations were made by Tan et *al*. while using the CRISPR-Cas3 systems for genome editing in eukaryotes: adding the Cas11 protein boosted the activity (Tan *et al*., 2022). Our previous data using the CRISPR interference tool (CRISPRi) show that in *Haloferax* CRISPRi is much more effective when the crRNAs used for CRISPRi are present in high concentrations (Stachler & Marchfelder, 2016). Thus, the concentration of crRNAs is an important factor for efficient Cascade activity. However, since a Cas11b free Cascade still binds and protects crRNAs effectively, the loss of interference is not due to lower crRNA concentrations.

McBride et *al*. showed for the bacterial I-D system, that Cascade without Cas11d does not bind to target DNA anymore. *E. coli* Cas11 helps to lock the nontarget strand by tightening the binding of Cascade to the DNA (Hayes *et al*., 2016), similar observations have been reported for other systems (Liu & Doudna, 2020; Xiao *et al*, 2017). Further experiments would have to reveal whether this is also true for the *Haloferax* Cas11b, being the reason why interference activity is reduced without Cas11b.

Taken together, the loss of interference activity might be due to inefficient interaction with the target DNA.

### Cas11b has a major impact on gene expression

Comparison of transcriptomes from cells with and without Cas11b show clear differences in expression of a plethora of genes. Absence of Cas11b results in down-regulation of more than a 100 hundred pHV4 genes in the region from HVO_A0279A-HVO_A0405. Since pHV4 is the element that encoded the CRISPR-Cas system, one can speculate that many genes within this element have coevolved with that system and their RNA stability depends on Cas11b to some extent. A few provirus genes are also affected by the absence ofCas11b, which might be related to its role in the CRISPR-Cas defence function. Since the function of many of the regulated genes (four sRNAs and 13 genes for hypothetical proteins) is not known, we cannot deduce the functional impact of Cas11b here. Genes involved in siderophore biosynthesis are upregulated in cells without Cas11b, thus the iron metabolism seems to be affected by a missing Cas11b, potentially by a de-stabilization of an inhibitory sRNA that has not been yet identified.

Taken together the many changes in gene expression upon loss of Cas11b suggest a role of this Cas protein beyond the original CRISPR-Cas function in defence and is hinting at a key role of Cas11b in several pathways.

## Data availability

Data from RNA-seq and mass spectrometry will be deposited in public databases and full supplementary data will be made available upon publication of the manuscript.

## Acknowledgments

We thank Manuela Weishaupt, Elena Katzowitsch and Panagiota Arampatzi for expert technical assistance and Lisa-Katharina Maier and Friedhelm Pfeiffer for constructive discussions and careful reading of the manuscript.

