## Supplementary material for "Internal in-frame translation generates Cas11b, which is important for effective interference in an archaeal CRISPR-Cas system": Suppl. Table1

Supplementary Figures

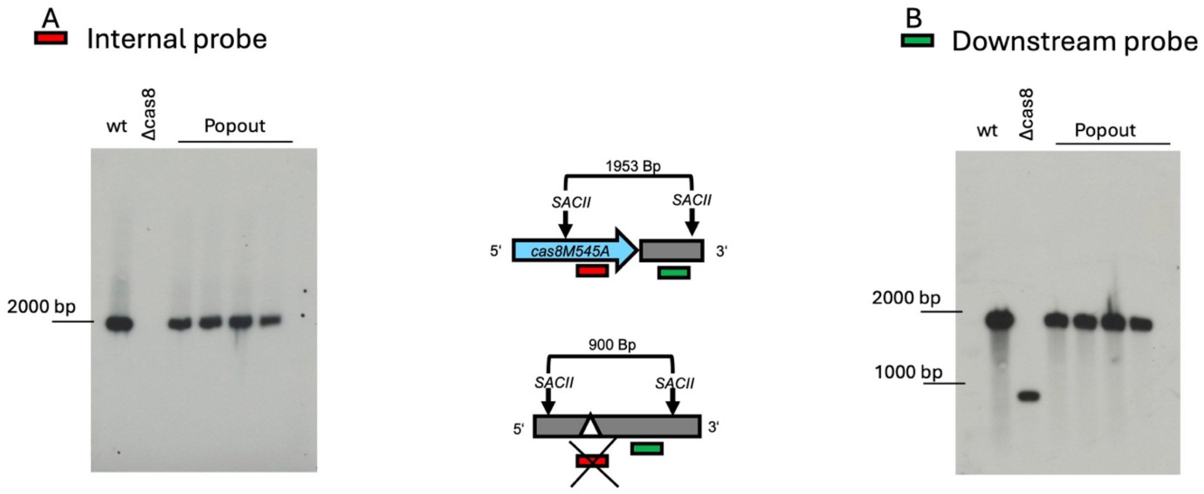

**Supplementary Figure 1. Conformation of strain *cas8*bM545A by Southern blot analysis.** To generate the mutant strain cas8M545A the mutated *cas8b* gene was re-integrated into the original genomic position of *cas8b* in a Δ*cas8b* strain. Four potential *cas8*M545A (HV120) clones were selected and checked for the presence of the mutated *cas8b* by southern blot analysis. To this end, *Sac*II digested gDNAs were separated on an 0.8 % agarose gel and transferred to a nylon membrane by capillary blot. The membrane was hybridised with radioactively labelled PCR probes. **A.** A probe against the *cas8*M545A gene was used. **B.** A probe binding to the downstream region of the gene was used. A schematic representation of the expected sizes of the fragments bound by the probes (red for the internal probe and green for the downstream probe) is shown in the middle. The corresponding autoradiographs are shown at the sides, sizes of the 1 kb ladder are indicated on the left of the films.

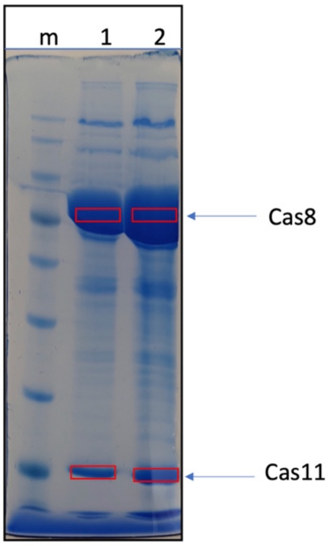

A.

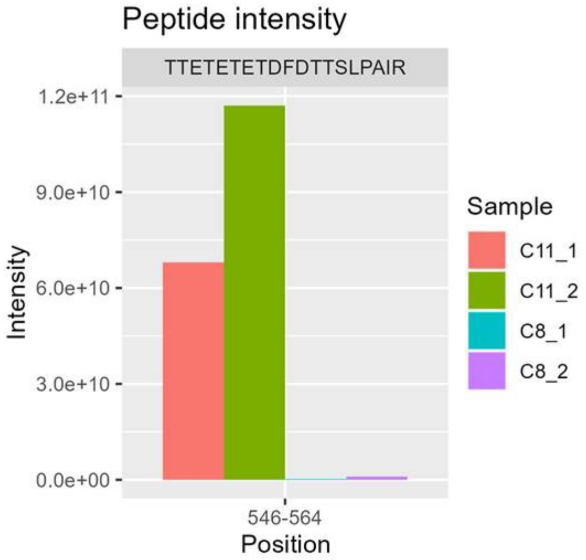

B.

**Supplementary Figure 2. Peptides identified for Cas11b. A.** Proteins identified as Cas8b and Cas11b in the western blot (Figure 1) were isolated from Coomassie stained SDS gels from two different lanes (red boxes) yielding samples C11_1, C11_2 for Cas11b, and C8_1 and C8_2 for Cas8b. **B.** If translation initiation starts at M545 the first peptide of Cas11b is MTTETETETDFDTTSLPAIR. Peptide TTETETETDFDTTSLPAIR was found with clearly higher intensity in the Cas11b samples, suggesting that Cas11b translation is initiated at M545. The N-terminal M is very often removed after translation in haloarchaea.

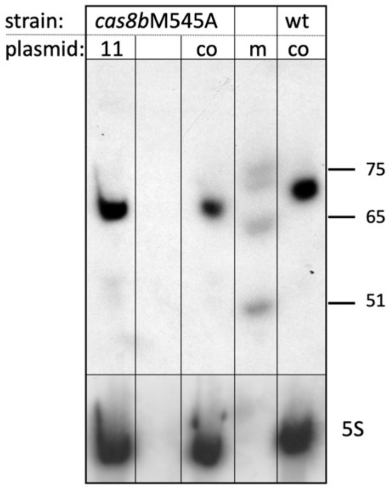

**Supplementary Figure 3. crRNA concentrations are not affected by Cas11b depletion.** In wild type *Haloferax* cells containing the *cas8b* gene on the chromosome (expressing Cas8b and Cas11b) crRNAs are readily visible (lane wt/ co; RNA from cells H119 x pTA927). The strain with the mutated *cas8b* gene (that does not express Cas11b) has similar amounts of crRNAs (lane *cas8*M545A/ co; RNAs from cells *cas8*M545A x pTA927). If the mutant strain is transformed with a plasmid expressing Cas11b again similar amounts of crRNAs are visible (lane *cas8*M545A/ 11; RNAs from cells *cas8*M545A x pTA927-*cas11b*). A DNA size standard is shown at the right, the upper panel shows hybridisation with a probe against a spacer of the P1 CRISPR locus, the lower panel shows hybridisation with a probe against the 5S rRNA.

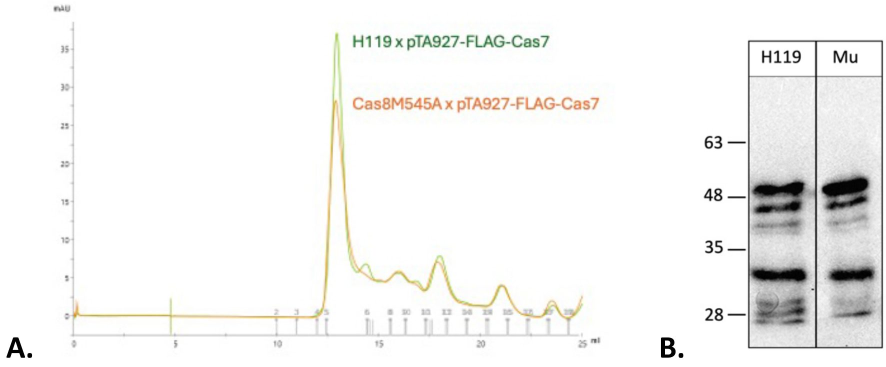

**Supplementary Figure 4. Western blot analysis of gel filtration fractions.** Cascade complexes from *Hfx. volcanii* can be purified via a FLAG-Cas7 co-purification with gel filtration as reported previously (Brendel, 2014). Cascade complexes from wild type cells (H119) and mutant cells *cas8*M545A, both transformed with a plasmid expressing FLAG-Cas7 were purified via FLAG-Cas7 using FLAG-agarose. Purified fractions were loaded onto a gel filtration column and fractions were collected. Proteins of fraction 6 from wild type cell and mutant cell extracts were loaded onto an SDS PAGE and subsequently transferred onto a western membrane, which was hybridised with an antibody against the FLAG tag. The full length Cas7-FLAG fusion protein is visible at about 48 kDa. Due to the high amount of acidic amino acids, halophilic proteins run generally slower on SDS PAGE, adjusted to this the calculated molecular weight is 47.4 kDa (Brendel, 2014; Guan *et al*, 2015), some degradation products of the Cas7-FLAG fusion protein are also detected .

**Supplementary Table 1. Strains, plasmids and primers used.**

**A. Strains**

| **Strains** | **Characteristics** | **Reference/ source** |
|---|---|---|
| *E. coli* DH5α | F- ϕ80d*lacZ*ΔM15 Δ(*lacZYA-argF*) U169 *deoR recA*1 *endA*1 *hsdR*17($r_k^-$, $m_k^+$) *gal-phoA supE*44λ- *thi*-1 *gyrA*96 *relA*1 | Invitrogen (Thermo Fischer Scientific, Waltham, MA, USA) |
| *E. coli* GM121 | F-, dam-3, dcm-6, ara-14, fhuA31, galK2, galT22, hdsR3, lacY1, leu-6, thi-1, thr-1, tsx-78 | (Allers *et al*, 2010) |
| H119 | DS70 (ΔpHV2), Δ*pyrE2*, Δ*trpA*, Δ*leuB* | (Allers *et al*, 2004) |
| Δ*cas8b* HV119 | DS70 (ΔpHV2), Δ*pyrE2*, Δ*trpA*, Δ*leuB*,Δ*cas8b* | (Stoll, 2013) |
| *cas8*M545A HV120 | DS70 (ΔpHV2), Δ*pyrE2*, Δ*trpA*, Δ*leuB, cas8b::cas8M545A* | this study |

**B. Plasmids**

| **Plasmids** | **Characteristics** | **Reference** |
|---|---|---|
| pTA131 | ColE1 ori, f1 ori, lacZ, $Amp^R$, pyrE2, | (Allers *et al.*, 2004) |
| pTA131-up.Cas8M545A.do | ColE1 ori, f1 ori, lacZ, $Amp^R$, pyrE2, upstream (UP) and downstream (DO) region of HVO_A0206 (UP: 546 bp + *Cas8bM545A* + DO: 501 bp) | this study |
| pTA131-up.do(Csh1) | ColE1 ori, f1 ori, lacZ, $Amp^R$, pyrE2, upstream (UP) and downstream (DO) region of HVO_A0206 (UP: 546 bp + DO: 501 bp) | (Cass *et al*, 2015) |
| pTA927-Cas7-N-Flag | ColE1 ori, f1 ori, lacZ, $Amp^R$, pHV2 ori, p.*tna*-promoter, L11e+terminator, *pyrE2*, *cas7*N-FLAG | (Stoll, 2013) |
| pTA927-FLAGcontrol | ColE1 ori, f1 ori, lacZ, $Amp^R$, pHV2 ori, pyrE2, L11e-terminator, p.*tna* -promoter, 3xFLAG, t.Syn-terminator | (Wörtz *et al*, 2022) |
| pTA927 | ColE1 ori, f1 ori, lacZ, $Amp^R$, pyrE2, pHV2 ori, p.*tnaA* promoter, t.syn terminator | (Allers *et al.*, 2010) |
| pTA927-FlagC | ColE1 ori, f1 ori, lacZ, $Amp^R$,pyrE2, pHV2 ori, p.*tnaA* promoter, 3XFLAG, t.syn terminator | (Hadjeras *et al*, 2023) |
| pTA927-*cas11b* | ColE1 ori, f1 ori, lacZ, $Amp^R$, pHV2 ori, p.tna-Promotor, L11e+Terminator, pyrE2, Cas11b, t.syn terminator | this study |
| pTA927-*cas8b*-FlagN | ColE1 ori, f1 ori, lacZ, $Amp^R$, pHV2 ori, p.*tna*-promoter, L11e+terminator, *pyrE2*, *cas8b*N-FLAG, t.syn terminator | (Cass *et al.*, 2015) |

| | | |
|---|---|---|
| pTA927-*cas8b*-FlagC | ColE1 ori, f1 ori, lacZ, $Amp^R$, pHV2 ori, p.*tna*-promoter, L11e+terminator, *pyrE2*, *cas8b*C-FLAG, t.syn terminator | this study |
| pTA927-*cas8b*M545A-FlagC | ColE1 ori, f1 ori, lacZ, $Amp^R$, pHV2 ori, p.*tna*-promoter, L11e+terminator, *pyrE2*, *cas8bM545A*-C-FLAG, t.syn terminator | this study |
| pTA927-*cas8bM545A*-FlagC-*cas11b* | ColE1 ori, f1 ori, lacZ, $Amp^R$, pHV2 ori, p.*tna*-promoter, L11e+terminator, *pyrE2*, *cas8b*C-FLAG, p.*tna*-promotor, *cas11b,* t.syn terminator | this study |
| pTA352 | ColE1 ori, f1 ori, lacZ, $Amp^R$, leuB, pHV1/4 ori | (Norais *et al*, 2007) |
| pTA352-PAM3-P1.1 | ColE1 ori, f1 ori, lacZ, $Amp^R$, leuB, pHV1/4 ori, PAM3 (TTC) followed by spacer 1 of CRISPR-locus P1) | (Maier *et al*, 2013) |

**C. Primers**

| **Oligonucleotide** | **Sequence** |
|---|---|
| Cas8_probe_do_rev | CGTCTTTATCGCTCGCCTCGAAGCTGAGC |
| Cas8_probe_intern_fwd | CGTGGGCCACCAGTTCACCGACTCG |
| Cas8_probe_do_fwd | TCCAACACATAACCAAACCAATGACGACACT |
| Cas8_probe_intern_rev | GGAACGATTCGAGTCTGTGTTCTCG |
| P1SP1 | GTTCCGGGAGGTCGCCGGTCGAGATGCCTGC |
| 5S | CGCAGGTGAGCTTAACTTCCGTGTTCGGG |
| 5-Csh1-NdeI | TATTATCATATGACAGGTCCAGATATCGACGACTTC |
| 3-Cas8-SmaI | TATTATCCCGGGGTTCGTGGTGCTCTCAGCGGGTTC |
| 8M545A | TTCCAATCGAACCACCAACCGCGACAACCGAAACTGAAA |
| 8M545Ar | TTTCAGTTTCGGTTGTCGCGGTTGGTGGTTCGATTGGAA |
| 5-Cas11-NdeI | TATTATCATATGACAACCGAAACTGAAACTGAAACTGA |
| 5-ptna-BamHI | TATTATGGATCCGCCCGTTCTCGTCGCGCTCTCGAAGCTGTT |
| 3-Cas11-XbaI | TATTATTCTAGATTAGTTCGTGGTGCTCTCAGCGGGTTCTT |
| IP cas8 rev | CAGTCACTCGCCCGTGGAAGCG |
| IP cas8 fw | TCCAACACATAACCAAACCAATGACGACACT |
| 3-cas8-rev | [phos]TTAGTTCGTGGTGCTCTCAGCGGGTT |
| 5-cas8-fw | [phos]ATGACAGGTCCAGATATCGACGACTTC |

**Supplementary Table 2. Genes up- or down regulated on pHV4 in a Cas11 less strain.** Genes encoded on pHV4 that are up-regulated with log2FC > 2 and down-regulated with log2FC < 2 are shown. A few genes (e.g. HVO_A0344, A0345, A0348, A0365, A0370, A0389, A0390A, A0405) occur multiple times. These genes are disrupted (by transposon targeting, frameshift or in-frame stop codon), subregions are represented as distinct annotations, resulting them to be independently analysed.

**A. Genes up regulated**

| gene | annotation | log2FC |
|---|---|---:|
| HVO_A0508 | 1,2-phenylacetyl-CoA epoxidase subunit B, paaB | 2,99 |
| HVO_A0509 | 1,2-phenylacetyl-CoA epoxidase subunit C, paaC | 2,85 |
| HVO_A0507 | 1,2-phenylacetyl-CoA epoxidase subunit A, paaA | 2,85 |
| HVO_A0510 | DUF59 family protein | 2,82 |
| HVO_A0535 | peptidase M24 family protein | 2,47 |
| HVO_A0505 | enoyl-CoA hydratase, fadA4 | 2,06 |

**B. Genes down regulated**

| gene | annotation | log2FC |
|---|---|---:|
| HVO_A0348A | hypothetical protein | -9,66 |
| HVO_A0317 | ArsR family transcription regulator | -9,55 |
| HVO_A0386 | N-methylhydantoinase (ATP-hydrolyzing) B | -8,81 |
| HVO_A0331 | D-galactonate dehydratase | -8,62 |
| HVO_A0320 | conserved hypothetical protein | -8,60 |
| HVO_A0351 | conserved hypothetical protein | -8,49 |
| HVO_A0334 | conserved hypothetical protein | -8,48 |
| HVO_A0308 | conserved hypothetical protein | -8,40 |
| HVO_A0388 | Lrp/AsnC family transcription regulator | -8,32 |
| HVO_A0393 | conserved hypothetical protein | -8,24 |
| HVO_A0318 | conserved hypothetical protein | -8,16 |
| HVO_A0315 | conserved hypothetical protein | -8,02 |
| HVO_A0307 | Lrp/AsnC family transcription regulator | -7,95 |
| HVO_A0346 | XerC/D-like integrase | -7,92 |
| HVO_A0350 | conserved hypothetical protein | -7,85 |
| HVO_A0380 | ABC-type transport system periplasmic substrate-binding protein (probable substrate dipeptide/oligopeptide) | -7,73 |
| HVO_A0362 | PQQ repeat protein | -7,70 |
| HVO_A0369 | CopG domain protein | -7,66 |
| HVO_A0279_A | transcription elongation factor TFS | -7,66 |
| HVO_A0316 | conserved hypothetical protein | -7,64 |
| HVO_A0385 | N-methylhydantoinase (ATP-hydrolyzing) A | -7,61 |
| HVO_A0397 | HTH domain protein | -7,55 |
| HVO_A0326 | beta-D-galactosidase | -7,51 |
| HVO_A0374 | conserved hypothetical protein | -7,49 |

| | | |
|---|---|---|
| HVO_A0329 | 2-dehydro-3-deoxy-phosphogluconate / 2-dehydro-3-deoxy-phosphogalactonate aldolase, bacterial-type | -7,43 |
| HVO_A0377 | hydantoin racemase | -7,38 |
| HVO_A0281 | ABC-type transport system ATP-binding protein (probable substrate sugar) | -7,35 |
| HVO_A0378 | N-methylhydantoinase (ATP-hydrolyzing) B | -7,27 |
| HVO_A0394 | HTH domain protein | -7,19 |
| HVO_A0401 | Fido domain protein | -7,14 |
| HVO_A0348 | ISH7-type transposase ISHvo15 (nonfunctional) | -7,11 |
| HVO_A0379 | N-methylhydantoinase (ATP-hydrolyzing) A | -7,10 |
| HVO_A0368 | RelE family protein | -7,04 |
| HVO_A0288 | probable oxidoreductase (short-chain dehydrogenase family) | -6,97 |
| HVO_A0333 | SprT family protein | -6,93 |
| HVO_A0360 | conserved hypothetical protein | -6,91 |
| HVO_A0376 | probable Xaa-Pro dipeptidase | -6,90 |
| HVO_A0365 | conserved hypothetical protein (nonfunctional) | -6,85 |
| HVO_A0291 | conserved hypothetical protein | -6,85 |
| HVO_A0280 | IclR family transcription regulator | -6,75 |
| HVO_A0335 | DUF1028 family protein | -6,67 |
| HVO_A0283 | ABC-type transport system periplasmic substrate-binding protein (probable substrate sugar) | -6,66 |
| HVO_A0314 | conserved hypothetical protein | -6,62 |
| HVO_A0347 | conserved hypothetical protein (nonfunctional) | -6,58 |
| HVO_A0332 | IclR family transcription regulator GacR | -6,56 |
| HVO_A0365 | conserved hypothetical protein (nonfunctional) | -6,45 |
| HVO_A0311 | probable halocin (homolog to halocin C8) | -6,43 |
| HVO_A0336 | ABC-type transport system ATP-binding protein (substrate D-galactose) | -6,30 |
| HVO_A0306 | pyridoxal phosphate-dependent aminotransferase | -6,22 |
| HVO_A0287 | homolog to mandelate racemase / homolog to muconate lactonizing enzyme | -6,22 |
| HVO_A0305 | methylmalonate-semialdehyde dehydrogenase | -6,18 |
| HVO_A0313 | cro/C1 family transcription regulator | -6,17 |
| HVO_A0289 | enamine/imine deaminase | -6,14 |
| HVO_A0312 | HTH domain protein | -6,14 |
| HVO_A0322 | ABC-type transport system permease protein | -6,11 |
| HVO_A0341 | amidase (hydantoinase/carbamoylase family) | -6,08 |
| HVO_A0339 | ABC-type transport system periplasmic substrate-binding protein (substrate D-galactose) | -6,08 |
| HVO_A0372 | beta-lactamase domain protein | -6,07 |
| HVO_A0396 | SWIM zinc finger domain protein | -6,03 |
| HVO_A0286 | DUF187 family protein | -6,03 |
| HVO_A0330 | D-galactose / L-arabinose dehydrogenase (NADP) | -6,01 |
| HVO_A0356 | conserved hypothetical protein (nonfunctional) | -5,98 |
| HVO_A0370 | conserved hypothetical protein (nonfunctional) | -5,95 |
| HVO_A0324 | conserved hypothetical protein | -5,90 |
| HVO_A0370 | conserved hypothetical protein (nonfunctional) | -5,87 |
| HVO_A0402 | conserved hypothetical protein | -5,86 |

| | | |
|---|---|---|
| HVO_A0387 | cupin 2 barrel domain protein | -5,82 |
| HVO_A0398 | conserved hypothetical protein | -5,75 |
| HVO_A0345 | transport protein (probable substrate cationic amino acids) (nonfunctional) | -5,75 |
| HVO_A0375 | transcription initiation factor TFB | -5,74 |
| HVO_A0395 | conserved hypothetical protein | -5,72 |
| HVO_A0392 | death-on-curing family protein | -5,69 |
| HVO_A0384 | ABC-type transport system ATP-binding protein (probable substrate dipeptide/oligopeptide) | -5,65 |
| HVO_A0358 | conserved hypothetical protein | -5,65 |
| HVO_A0323 | ABC-type transport system ATP-binding protein | -5,62 |
| HVO_A0328 | 2-keto-3-deoxygalactonate kinase | -5,52 |
| HVO_A0361 | conserved hypothetical protein | -5,51 |
| HVO_A0381 | ABC-type transport system permease protein (probable substrate dipeptide/oligopeptide) | -5,50 |
| HVO_A0338 | ABC-type transport system permease protein (substrate D-galactose) | -5,49 |
| HVO_A0344 | UspA domain protein (nonfunctional) | -5,42 |
| HVO_A0353 | ISH3-type transposase ISH51 (nonfunctional) | -5,42 |
| HVO_A0370 | conserved hypothetical protein (nonfunctional) | -5,26 |
| HVO_A0296 | probable oxidoreductase (short-chain dehydrogenase family) | -5,23 |
| HVO_A0383 | ABC-type transport system ATP-binding protein (probable substrate dipeptide/oligopeptide) | -5,20 |
| HVO_A0399 | conserved hypothetical protein | -5,13 |
| HVO_A0342 | IclR family transcription regulator | -4,95 |
| HVO_A0389 | HTH domain protein (nonfunctional) | -4,94 |
| HVO_A0400 | conserved hypothetical protein | -4,93 |
| HVO_A0344 | UspA domain protein (nonfunctional) | -4,91 |
| HVO_A0403 | ISHwa16-type transposase ISHvo16 | -4,91 |
| HVO_A0371 | conserved hypothetical protein | -4,84 |
| HVO_A0310 | conserved hypothetical protein | -4,78 |
| HVO_A0366 | conserved hypothetical protein | -4,76 |
| HVO_A0295 | amidase (hydantoinase/carbamoylase family) | -4,68 |
| HVO_A0345 | transport protein (probable substrate cationic amino acids) (nonfunctional) | -4,63 |
| HVO_A0382 | ABC-type transport system permease protein (probable substrate dipeptide/oligopeptide) | -4,60 |
| HVO_A0282 | creatininase domain protein | -4,53 |
| HVO_A0303 | probable allantoinase | -4,42 |
| HVO_A0390_A | conserved hypothetical protein (nonfunctional) | -4,42 |
| HVO_A0299 | ABC-type transport system periplasmic substrate-binding protein | -4,35 |
| HVO_A0321 | conserved hypothetical protein | -4,35 |
| HVO_A0357 | conserved hypothetical protein | -4,35 |
| HVO_A0290A | 2-keto-3-deoxygluconate kinase (nonfunctional) | -4,32 |
| HVO_A0284 | ABC-type transport system permease protein (probable substrate sugar) | -4,26 |
| HVO_A0337 | ABC-type transport system permease protein (substrate D-galactose) | -4,26 |
| HVO_A0290 | 2-dehydro-3-deoxy-phosphogluconate aldolase, bacterial-type | -4,25 |
| HVO_A0297 | ABC-type transport system permease protein | -4,21 |

| | | |
|---|---|---|
| HVO_A0389 | HTH domain protein (nonfunctional) | -4,12 |
| HVO_A0319 | hypothetical protein | -4,07 |
| HVO_A0327 | conserved hypothetical protein | -3,96 |
| HVO_A0348 | ISH7-type transposase ISHvo15 (nonfunctional) | -3,69 |
| HVO_A0294 | ABC-type transport system ATP-binding protein | -3,58 |
| HVO_A0390_A | conserved hypothetical protein (nonfunctional) | -3,54 |
| HVO_A0301 | probable polysaccharide deacetylase | -3,53 |
| HVO_A0391 | conserved hypothetical protein | -3,42 |
| HVO_A0405 | conserved hypothetical protein (nonfunctional) | -3,37 |
| HVO_A0293 | ABC-type transport system ATP-binding protein | -3,33 |
| HVO_A0292 | transport protein (probable substrate cationic amino acids) | -3,27 |
| HVO_A0405 | conserved hypothetical protein (nonfunctional) | -3,19 |
| HVO_A0325 | UPF0121 family protein | -2,91 |
| HVO_A0298 | probable oxidoreductase (short-chain dehydrogenase family) | -2,88 |
| HVO_A0389 | HTH domain protein (nonfunctional) | -2,66 |
| HVO_A0302 | asparaginase/glutaminase family protein | -2,63 |
| HVO_A0285 | ABC-type transport system permease protein (probable substrate sugar) | -2,62 |
| HVO_A0295_A | luciferase family protein | -2,53 |
| HVO_A0023 | DUF1931 domain protein | -2,48 |
| HVO_A0359 | conserved hypothetical protein (nonfunctional) | -2,47 |
| HVO_A0300 | ABC-type transport system permease protein | -2,47 |
| HVO_A0363 | conserved hypothetical protein | -2,45 |
| HVO_A0309 | ISH7-type transposase HfIRS6 (nonfunctional) | -2,40 |
| HVO_A0359A | conserved hypothetical protein (nonfunctional) | -2,01 |
